# TOP-DOWN CLIMATIC PROCESSES MODULATE BIODIVERSITY-FUNCTIONING RELATIONSHIPS ACROSS NORTH AMERICAN FORESTS

**DOI:** 10.1101/2024.11.04.621856

**Authors:** Xavier Serra Maluquer, Oscar Godoy, Paloma Ruiz-Benito, Julen Astigarraga, Francisco Rodríguez-Sánchez, Julián Tijerín-Triviño, Marta Rueda

## Abstract

Experimental evidence indicates that more diverse communities sustain higher levels of ecosystem functioning. Generally, bottom-up processes accounting for community selection and complementarity effects on biodiversity are used to explain biodiversity effects on ecosystem functioning. However, top-down macroecological processes can also influence the distribution of ecosystem functions across large environmental gradients and, importantly, biodiversity patterns. Here, we tested whether past climate instability, current climate, and climate-driven species abundance explain tree richness gradients, and therefore, determine the relationship between species richness and ecosystem functioning using 137,808 forest inventory plots across continental extents including forests of Canada, the US, and Mexico. Consistent with previous research, we found a positive monotonic relationship between species richness and biomass (measured as level stand basal area) across North American forests. However, structural equation models revealed that the indirect effect of past climate instability, current climate conditions, and climate-driven individuals’ abundance underlie species richness patterns and the positive covariation between species richness and biomass. The importance of top-down climatic processes varied from boreal to tropical regions, yet their significant effects were pervasive across all forest types, indicating that climate strongly modulates the strength of the relationships between species richness and basal area. Our results help to understand the assembly processes driving the ubiquitous positive monotonic relationship between species richness and basal area across continental extents. This knowledge, obtained from adopting a macroecological perspective, helps understanding biodiversity-functioning relationships, which are usually explained by mechanisms at the community level.

**Significance statement:** More biodiverse communities sustain higher levels of ecosystem functions (BEF hypothesis). The role of biotic interactions among trees within ecological communities determining BEF has been widely studied, but it is not well known how past climatic variability, current climate, and species abundance determine BEF. Using 137,808 forest inventory plots across North American forests located in Canada, the US, and Mexico, we document a positive relationship between tree species richness and stand basal area. Detailed analysis revealed that the positive BEF relationship is underlined by a strong and indirect effect of past and current climate. Our results highlight the importance of top-down climate processes operating at macroecological scales to understand biodiversity-functioning relationships.

## Introduction

During the last decades, a substantial body of experimental evidence has demonstrated that more diverse communities sustain higher levels of functioning than less diverse ones (1–3). Explanations for this positive and stabilizing effect of biodiversity on ecosystem functioning (hereafter BEF) commonly involve bottom-up processes operating at local scales, such as selection effects (i.e. high performance through competitive dominance) and complementarity effects (i.e. niche partitioning and facilitation) (4). BEF research has predominantly focused on the critical role of biodiversity, with extrapolations of empirical findings to natural ecosystems to understand how changes in richness and functional composition of communities influence functioning across space (5, 6). However, to understand and predict BEF relationships across large geographic areas, it is paramount to additionally consider the effect of variables underlying diversity patterns and ecosystem functioning through top-down processes (7–10). Thus, the shape and strength of BEF relationships can be strongly modulated by climate, soil conditions, human activity, and biogeographical history among others (11–18). However, it is not well known within BEF research how these top-down processes influence biodiversity and, therefore, the magnitude of BEF relationships. In fact, it is critical to understand to what extent BEF patterns and their underlying mechanisms are universal due to the recognized bottom-up processes or if they are environmentally dependent due to top-down processes, particularly as we study larger geographic areas from local communities to continents. Explicitly including how top-down processes determine biodiversity, and hence its relationship with ecosystem function, will enable us to describe the generality of BEF across spatial scales, despite the possibility of being driven by different processes.

North American forests provide a unique opportunity to test top-down processes in driving BEF relationships. If all North American forests were similar, we would expect that local processes would govern BEF relationships. Remarkably, the reality is different. North American forests encompass a diverse array of forest types, ranging from semitropical to boreal forests, distributed continuously along both latitudinal (from Yucatan to Alaska) and longitudinal (from California to Quebec) gradients. Given the considerable variation in species richness and biomass stocks across these forests (19, 20), we hypothesize that top-down processes will generally exert a significant influence on the continental-scale relationship between biodiversity and biomass. More specifically, we expect this relationship to depend on three primary hypotheses, not mutually exclusive, that account for a series of climatic, ecological, and evolutionary drivers on species richness and, therefore, species richness vs. biomass relationships (see Fig. 1). First, past climate can modulate species diversity (21), with regions that have experienced greater climate stability over the past millennia hosting higher numbers of species and phylogenetic diversity than regions subjected to species extinctions due to pronounced climatic fluctuations (21, 22) (**H1: past climate instability hypothesis**, Fig. 1). For example, temperate tree flora shows higher phylogenetic clustering in regions where climatic extinctions have been documented due to past climatic events (23). Second, the diversity of tree species in North America forests follows a gradient of temperature and precipitation (24), with wet and warm sites hosting more species at macroecological scales (8). This positive current climate-diversity pattern may stem from a combination of evolutionary and ecological hypotheses. Areas with higher temperatures and more precipitation result in higher diversity because it may lead to higher speciation rates and lower extinction (**H2: current climate hypothesis**, Fig. 1) (25), as well as larger and more viable population sizes (26, 27) (**H3: climate-driven abundance hypothesis**, Fig. 1). We further hypothesize that the three proposed hypotheses are likely to have differential effects on species richness variability across different forest types of North America, influenced by the water-energy dynamics and the extent of past climate change experienced within each forest type. Therefore, testing their effects across different forest types will help to understand the BEF context dependency. Furthermore, these three different hypotheses do not deny the fact that other biotic factors may also play a role in shaping BEF relationships in North American forests. Other events determining biomass stock and productivity might not be as deterministic as environmental conditions or have such large-scale continental variation, but they can be even more important than environmental conditions in modulating BEF relationships. These are the cases of forest structure and disturbance events such as fire, hurricane, and logging activities (28).

**Figure 1.**
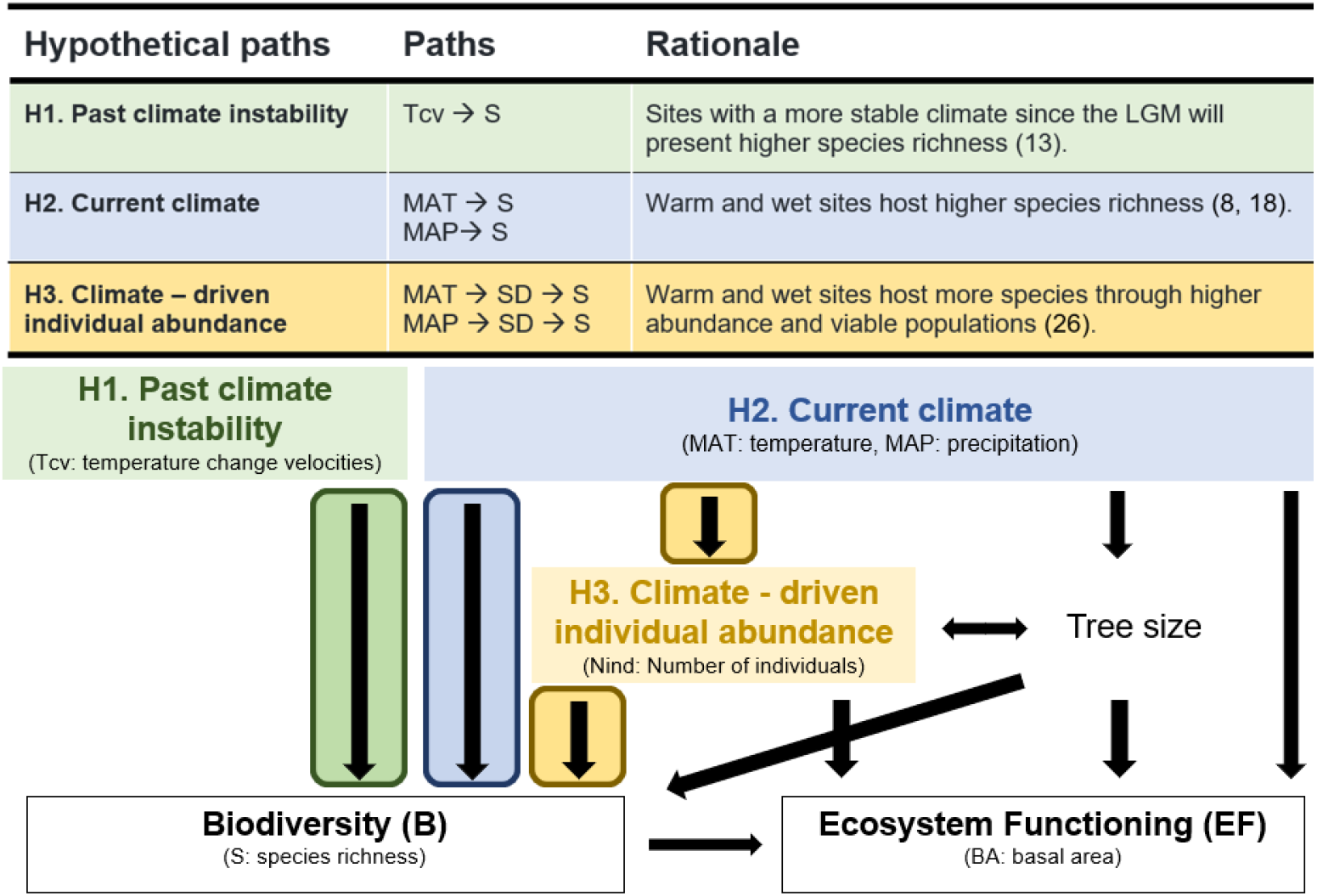
Hypothesis underlying the three proposed macroecological top-down effects of climate on biodiversity and biodiversity-ecosystem functioning relationship. We provide a table and conceptual path model indicating our three hypotheses of top-down processes (i.e. past climate instability, current climatic conditions and climate-driven individual abundance) driving biodiversity and BEF (Biodiversity - Ecosystem Functioning, see white boxes with black border). The effect of each of the three climatic hypotheses on biodiversity are shown with arrows, and highlighted in colors for clarity. See Appendix A of Supplementary Material for a detailed description and justification of all the included paths.

To what extent integrating top-down macroecological hypotheses of species distribution and diversity can improve our understanding of BEF relationships across continental areas, such as North American forests, has not been previously explored. Here, we combine data from four national forest inventories, encompassing over 137,808 plots from boreal to tropical forests. We integrate this data with information of past climate instability, current temperature and precipitation, two well-known drivers of forest productivity (29), and forest structural properties including individual abundance and maximum tree size. We employ structural equation models to test the three specific macroecological hypotheses that might influence species richness and, therefore, the biodiversity-biomass relationships that emerge at continental level (Fig. 1). Specifically, we expect a positive relationship between species richness and basal area (a good proxy of forest biomass, (30, 31)). We hypothesize that a positive BEF relationship will exhibit a more linear trend in forest types with limiting environmental conditions for growth, such as water- or temperature-limited forests. In contrast, we expect saturating relationships to occur in areas with more favorable conditions, such as tropical or temperate forests, which tend to have higher species richness and greater growth potential due to favorable environmental conditions(15).

## Results and discussion

We found a positive monotonic relationship between species richness and biomass across North American forests, measured as stand level basal area (Fig. 2A). This relationship aligns with previous studies reporting positive BEF relationships in forests worldwide, with an estimate value of 0.35 (previous work found a mean estimate of 0.26, SD = 0.09) (6). We also found that our three top-down hypotheses operating at the macroecological scale largely influence these BEF relationships. Regarding past climate instability, measured as temperature change velocities (H1), we observed a stronger BEF relationship with more positive exponents, that is, with more change in temperature (Fig. 2B). These results imply that areas of North America with less climatic stability since the last Glacial Maximum (approximately 20,000 years ago) exhibit stronger BEF relationships. In general, regions characterized by high velocities in temperature change correspond to those once covered by the Laurentide ice sheet (32) and hold fewer species. Because of this limitation in species richness, these forests are restricted to the steepest, and therefore strongest, part of the BEF relationship between species richness and basal area. An example of such forest areas in our dataset are the boreal forests, primarily represented by the Quebec forest inventory (blue points in Fig. 2A), for which previous work has also demonstrated stronger BEF relationships compared to temperate forests (15). For current climate conditions (H2), our results indicate stronger BEF relationships in regions with lower temperatures (Fig. 2B) but higher precipitation (Fig. 2C, D). The combination of both climatic variables underscores the importance of aridity. While previous studies have predicted and documented the effect of aridity on reducing basal area (30, 33), its role in modulating BEF at continental scales has only been described for drylands (12). However, our results suggest that aridity also plays a significant role in modulating BEF across continental forests. Finally, we found that the number of individuals (H3) also has a strong effect on the relationship between species richness and basal area (Fig. 2D). This result indicates the paramount importance of accounting for tree density when evaluating BEF relationships. The role of abundance in BEF can be viewed from two alternative perspectives. First, comparing forest plots with similar species richness levels, reveals that those with more individuals can attain more basal area. Yet, abundance can also influence species’ richness through two complementary ways. The most parsimonious option is the occurrence of a sampling effect (34, 35), that is, an increase from two to three individuals raises the probability of increasing from one to two species. Complementarily, ecological limitations may also occur according to the more individuals hypothesis (26), with harsh environments, such as cold or arid forest plots, limiting the occurrence of more species due to the impossibility of sustaining species with enough population sizes. In sum, our results indicate that the observed monotonic relationship between basal area and species richness is simultaneously modulated by top-down climatic drivers.

**Figure 2.**
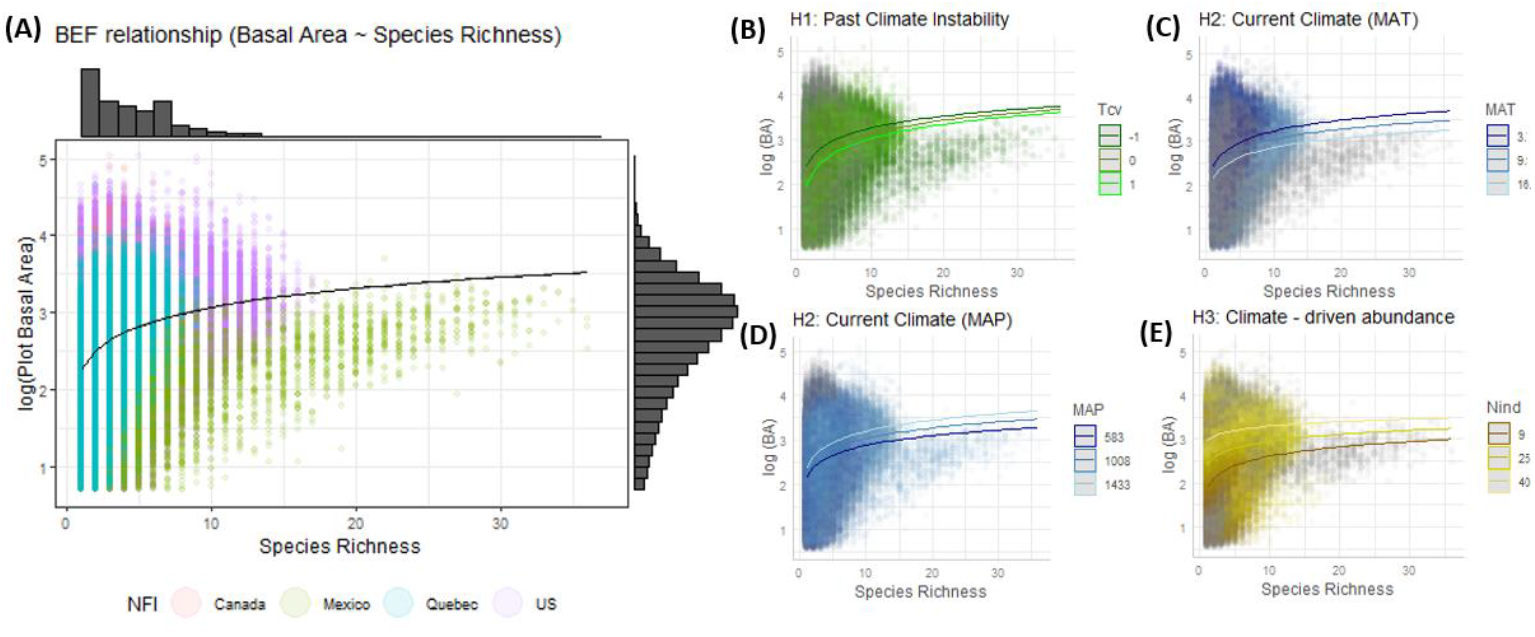
Relationships between diversity and ecosystem functioning for North American forests and its variation depending on past climate, current climate and climate-driven individual abundance. (A) Relationship between species richness and basal area across North American forests. The black line shows the predicted log basal area in US plots of common area (accounting for plot area). Predictions for each inventory can be seen in Fig. SI 1. (B-E) Relationships between basal area and species richness depending on: (B) past climate (i.e. temperature change velocities, km yr^−1^); (C, D) current climatic conditions (i.e. temperature and precipitation, °C and mm); and (E) climate-driven individual abundance (i.e. number of individuals, No. trees, predictions rounded) showing a predicted line and values for 10, 50 and 90 percentiles.

Given that past climate instability (H1), current climate conditions (H2) and climate-driven abundance (H3) modulate BEF relationships between species richness and basal area, we performed structural equation models (SEMs, see methods and SI for details) following our a priori causal relationships (Fig. 1) to explore the effects of these three top-down climate processes on species richness, and on the relationship of species richness with ecosystem functioning. We found a positive and direct, although weak, effect of diversity on ecosystem functioning (standardized coefficient 0.02, Figure 3). Remarkably, we observed a greater effect of top-down climate processes on both species richness and stand basal area across North American forests (Fig 3). The consistent effect for all variables (range of effects between 0.13-0.29 as shown in the standard coefficients, see Fig 3), indicates the non-exclusive importance of past climate instability, current climate conditions, and abundance in explaining patterns of tree species richness and BEF relationships at continental scale, even after accounting for other structural variables at the forest plot level, such as tree size, known to significantly influence BEF relationships (36).

**Figure 3.**
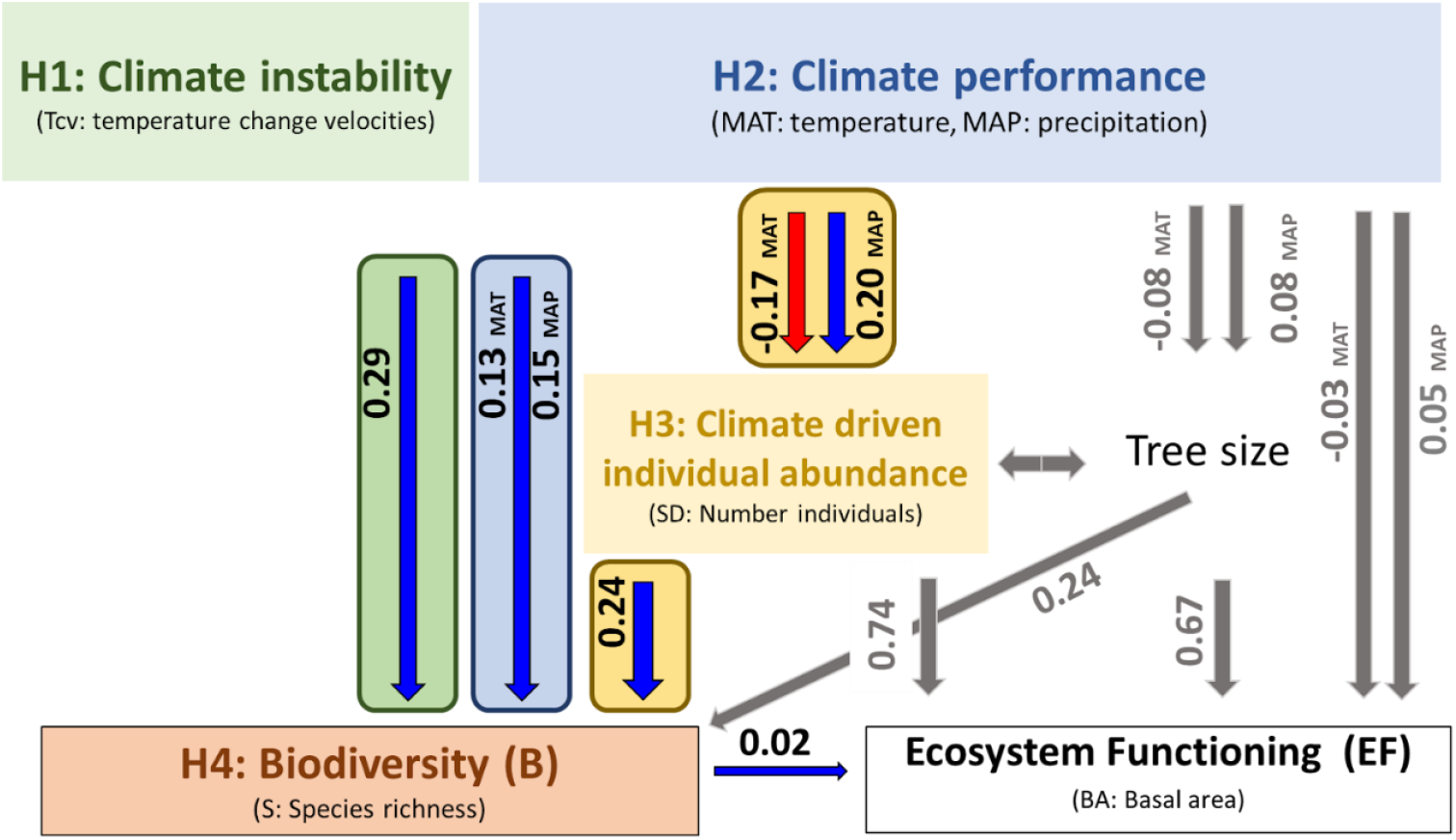
Evidence for the macroecological climatic hypothesis driving diversity and, therefore, diversity-ecosystem functioning in North American forests. We show the structural equation model results showing standardized coefficients next to arrows. Blue arrows represent positive relationships and red arrows negative relationships, whereas in gray are shown the relationships that were not specific hypotheses (see Fig. 1). For arrows coming from the climate box, whether they represent MAP or MAT is indicated. Correlations between exogenous variables (past climate instability and current climate) are shown in Fig S3.

Despite we found support for the three climatic hypothesis driving continental patterns of tree species richness, the direction of some relationships was unexpected. Firstly, for the first hypothesis and contrary to our expectations, we observed a positive effect of temperature change velocities on species richness (standard coefficient of 0.29, testing H1 in Fig. 3). This result suggests that regions that have been more climatically stable (i.e. lower temperature change velocities) since the Last Glacial Maximum had a lower number of tree species. This counterintuitive result can be explained by historical differences between the temperate forest of the western and eastern parts of North America (see SI Fig. S4). Currently, temperate forests of the east coast harbor greater tree diversity than those on the Pacific slope (21), despite being in areas with higher temperature change velocities due to the expansion and retraction of ice sheets and greater topographical complexity (SI Appendix, Fig. S4). For the second hypothesis, we observed a direct positive effect of current temperatures (standard coefficient of 0.13, testing H2 in Figure 3) and of precipitation (standard coefficient of 0.15, testing H2 in Figure 3) on tree species richness, as expected. This finding agrees with the idea that warm and wet sites sustain more species compared to cold and arid ones (8). Finally, regarding the third hypothesis, we found that regions with more individuals support a greater number of species, as expected (26). Specifically, higher viable abundances lead to higher numbers of species (standard coefficient of 0.24, testing H3 in Fig. 3). The indirect effects of both temperature and precipitation on biodiversity through abundance (standard coefficients of −0.17 and 0.2 effect on abundance, respectively, H3 in Fig. 3) indicate that sites with relatively low temperatures and high precipitation promote species richness by increasing species abundances. In other words, sites with high temperatures and low precipitation, such as dry ecosystems, harbor lower numbers of individuals, likely belonging to lower number of species.

According with the wide variety of forest types across North America, we found widespread variation in the strength and sign of the causal structure of direct and indirect effects previously reported (Fig. 4). Most notably, we detected changes in the sign of the relationship between past climate instability (H1) and species richness. Forest types with relatively stable climate, such as mountain and tropical forests, exhibited a positive effect of past climate instability on species richness, whereas forest types with greater past climate instability, such as boreal, temperate continental and subtropical humid forests, showed the opposite pattern (Fig. 4A & F). These results suggest that the expected relationship of H1 is confirmed for forest types with large divergences in past climate instability such as boreal ones. This is particularly evident in eastern North America, which experienced the highest gradient in climate change velocities and changes in ice sheet extent and has less topographical complexity (Fig. S4), and where extinctions due to climatic changes have been more pronounced. In addition, the effect of current climate (H2) varied from a positive effect of mean temperature on species richness in the northeastern forests (i.e. from temperate to boreal forests) to a negative effect in tropical and subtropical forests (Fig. 4B). This result agrees with the importance of energy and water limitations, as well as trait adaptation in shaping continental gradients of species richness across forest types (8). When energy is limiting, the tropical niche conservatism hypothesis posits that a few taxonomic tree groups have evolved traits allowing them to cope with harsh cold climates as species move northward, while warmer regions support a wide diversity of tree species with disparate functional forms and evolutionary backgrounds (37, 38). When significant, mean annual precipitation (H2) had a positive effect on species richness. The strongest magnitude of this effect was found in subtropical mountain forests, where driest conditions are found (Fig. 4C & H). Conversely to this variation across forest types, we also observed some consistent relationships. Species abundances (H3), measured as the number of individuals, were positively associated with species richness (H3) across all forest types. Finally, we found a positive direct effect of species richness on stand basal area, with a similar BEF relationship in temperate mountain, continental and tropical forests.

**Figure 4.**
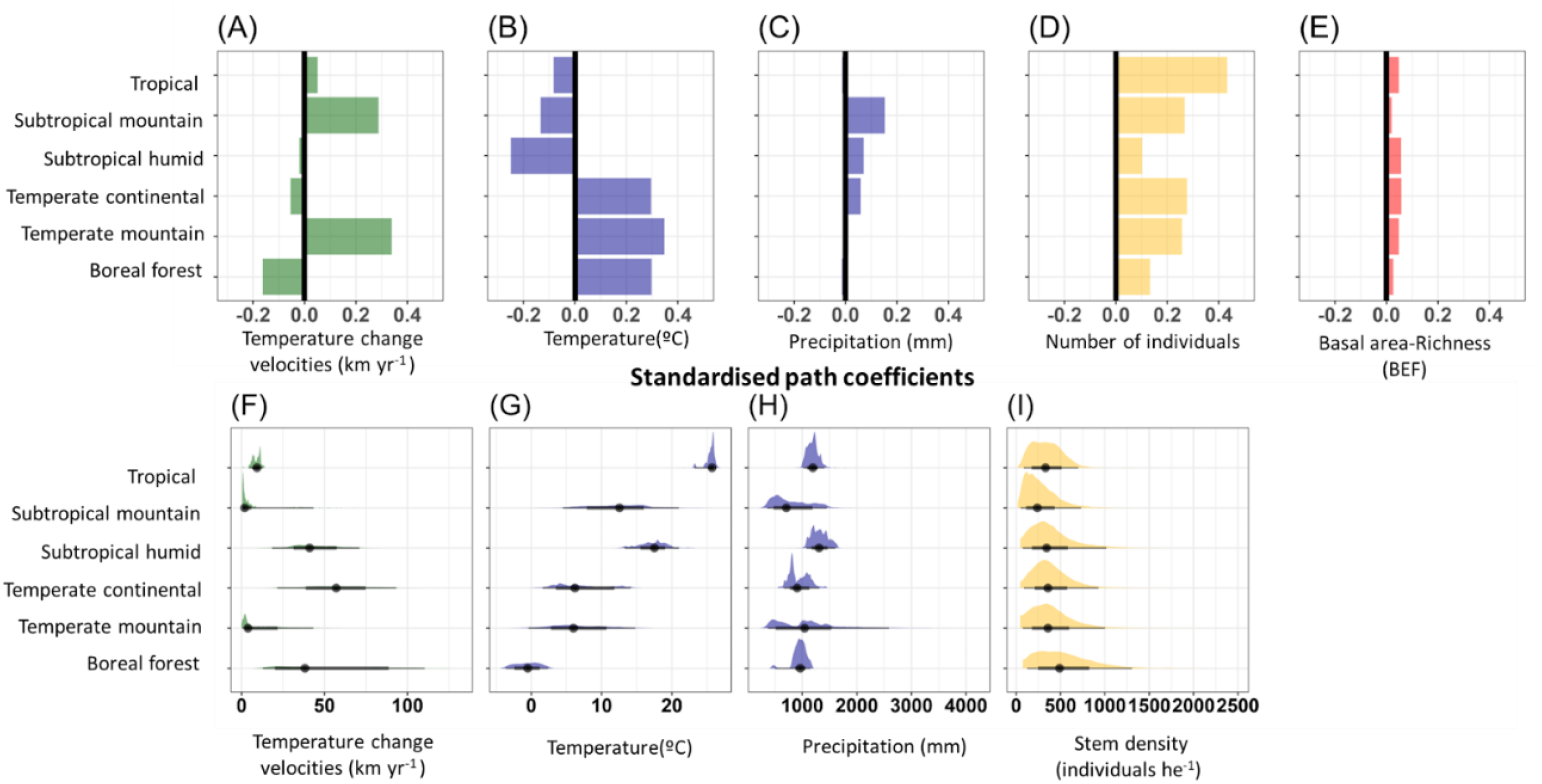
Direct effects of macroecological top-down climatic processes driving biodiversity and richness-diversity relationships across different forest types. The upper panels show standardized path coefficients of direct effects of temperature change velocities (H1. Past climate instability), temperature and precipitation (H2. Current climate hypothesis), and number of individuals (H3. Climate-driven abundance) on species richness and biomass across forest types. The lower panels show distribution of temperature change velocities (km yr^−1^), mean annual temperature (°C), annual precipitation (mm) and stem density (No. individuals ha^−1^, standardized by area to make it comparable between inventories) across forest types.

Taken together, we report a positive monotonic relationship between tree species richness and basal area across North American forests, indicating that their BEF relationships are consistent with those observed in other forests and ecosystems worldwide (6, 12). However, our findings present novel insights into how multiple facets of past and present climate indirectly influence BEF through changes in species richness. While we overall find robust support for the top-down effects of the three climate hypotheses studied (past climate instability (H1), current climate conditions (H2) and climate-driven abundance (H3)), detailed analyses also reveal marked differences across forest types. Specifically, in eastern forests, where there have been higher divergences in climate over the last 20,000 years, climate instability leads to a strong negative effect of species richness.

Meanwhile, current climatic controls (mean annual temperature) have differing effects on BEF, depending on whether forests are cold- or water-limited. Finally, the effect of climate-driven abundance indirectly modifying species richness is especially strong in tropical forests. This finding indicates that the number of individuals most notably increases species richness in hyperdiverse systems like tropical forests. Both the general top-down climate effects as well as their changes in relative importance across forest types suggest that different mechanisms operate at different spatial scales. At the continental level, the indirect effects of climate arise from the aggregation of forest types with different flora, functional characteristics, and evolutionary histories. Meanwhile, results at the regional scale (i.e., within forest types) suggests that ecological processes limit species establishment and abundance. In sum, our results do not deny the positive effect of biodiversity on functioning. Instead, they provide compelling evidence that integrating macroecological hypotheses on species distributions complements local scale insights of bottom-up processes, thereby deepening our understanding of continental patterns in the relationship between biodiversity and ecosystem functioning.

## Material and Methods

### Forest Inventory Data

To test the effect of the macroecological hypotheses on the relationship between species richness and stand basal area at the continental level (Fig. 1) we compiled data from the four forest inventories spanning North America, including Mexico, the United States, Quebec and Canada, (SI Appendix Fig. S1). We restricted the data to forest plots measured within the same decades (from 2000 to 2016).

For Mexico we used the second National Forest and Soil Inventory (INFyS, 2009-2014). INFyS employs a stratified sampling design, covering approximately 26,000 plots across Mexico’s territory, with a sampling intensity of 5 - 10 km^2^ depending on forest type. Plot design is as follows: one hectare plot comprising four subplots of 400 m^2^ each. In subplots, all trees with a diameter at breast height (d.b.h.) greater than 7.5 cm are measured. From all sampled plots we included those that meet the following criteria: 1) plots classified as tropical forests or forests, excluding those categorized as other communities such as shrublands and arid communities; 2) plots where disturbance impact (including fire, pests, wind, human, etc.) is not perceptible or minor; 3) plots with woody species alive (i.e., non woody species, such as Cactaceae and arborescent Pteridophyta were excluded) and with taxonomic information provided at least at genus level.

For the United States of America (US), we used the Forest Inventory Analysis (FIA) database. FIA plots cover all forested areas of the US (and portions of Alaska). Since 2000, FIA follows an annual inventory sampling where part of the plots (10-20%) of all states (except Alaska) are re-measured. Circular plots of 0.4 hectares are used, with four subplots of 24-feet radius (7.3 m, 0.0167 ha), where the diameter and height of all trees greater than 12.7 cm d.b.h. is recorded. We selected plots with the following criteria: 1) classified as non-disturbed forest; 2) first census after 2000.

For Quebec we used the 4th Inventoire Ecoforestiere of Placettes Énchantillons Permanents (PEP), which includes approximately 12,000 plots sampled between 2003 and 2016. Plots have a stratified design with an aggregated radius of 11,28 m (400 m2) and 14 m, where different sized trees are measured. For this study, we only kept data from the 11,28 radius, where all trees larger than 9 cm of d.b.h are measured, with tree size and species identity recorded. In our analyses, we selected plots that had no signs of disturbance in more than the 25 % of their area.

For Canada, we used the Canada National Forest Inventory (NFI). The Canadian NFI includes a small set of approximately 1,114 permanent ground plots established between 2000 and 2006 and re-measured between 2008 and 2017. Plots are circular (although with different sizes) where all trees with d.b.h. > 9 cm are sampled. We selected plots using the variables land base and land cover, which indicate if the plots are vegetated and contain tree.

We calculated stand diversity and structural variables using only alive trees from the four forest inventories using census from 2000 to 2016, with a criterion of at least three trees present in the plot. We used a total of 137,808 plots (Mexico = 6,815, US = 121,384 Canada = 700, and Quebec = 8,909). To ensure consistency across datasets, we focused on trees with a d.b.h. > 12.7 cm, aligning with the threshold used in the FIA dataset. Tree species richness was calculated as the total number of tree species present in each plot (No. tree species); stand abundance was determined as the number of alive trees present in each plot (No. trees); maximum tree size was defined as the 95th percentile of all trees present in each plot (cm); and stand basal area was calculated as the sum of the individual basal area per plot (m^2^ ha^−1^). Plots that had a basal area lower than two m^2^ ha^−1^ or three individuals were discarded.

### Climatic Information

Annual precipitation and annual mean temperature were gathered for each plot from Worldclim v2 (39). We also obtained temperature change velocities from the late Quaternary to the present (40). These climate change velocities (km yr^−1^) represent the spatial rate of temperature changes between the last glacial maximum (around 20,000 years ago) to the present. Previous research has shown that velocities in temperature changes influence the number of species as well as ecosystem functions (13, 40). Because climate change velocities are directly linked to topographic variation, smaller velocities are expected in mountain areas, whereas much larger velocities occur in flat areas (40). Maps of the mean annual temperature, mean annual accumulated precipitation and temperature change velocities can be found in SI *Appendix* Fig. S4.

### Statistical analyses

#### Relationship between diversity and biomass

To assess the Biodiversity-Ecosystem-Functioning (BEF) relationship between tree species richness and standing basal area, we first modeled log-transformed basal area assuming a Gaussian distribution as a function of log-transformed species richness, including plot area as a covariate to account for National Forest Inventories (NFIs) sampling differences. In addition to evaluate how the relationship between species richness and basal area is influenced by hypotheses H1, H2 and H3 independently, we fitted three additional models (with log-transformed basal area, assuming Gaussian distribution): the first model included temperature change velocities, the second included mean annual temperature and annual precipitation, and the third one included number of individuals. In all cases, these models incorporated the single interaction between each of the included variables and species richness.

#### Structural Equation Model (SEM)

Based on a priori knowledge and aiming to discern the importance of the three proposed macroecological hypotheses (Fig. 1, See SI *Appendix* A Further details on Structural Equation Model) on the continental-scale relationship between species richness and stand basal area, we constructed a structural equation model (SEM) (41). The SEM contains the direct relationship between past and present climate with species richness and stand structure (number of individuals and maximum tree size) as well as the indirect effects of climate on species richness and basal area through stand structure (number of individuals and maximum tree size).

We fitted the theoretical SEM model, based on prior knowledge available in the literature (Fig. 1, SI Appendix A for a detailed description of path hypotheses, model parameterization and evaluation), using data from the four national forest inventories (Mexico, US, Quebec and Canada). We used a piecewise SEM approach (42), which allows different model structures between the relationships. We employed the “piecewiseSEM” package (43) in R (44). Piecewise SEM allows fitting each equation of the SEM separately, rather than evaluating the whole variance-covariance matrix. This procedure is required when data analyses need flexibility in terms of selecting different data distributions and hierarchical structure (42).

After fitting the SEM using data for all North American forests, we fitted separate models for six different forest types: boreal, temperate mountain, temperate continental, subtropical humid, subtropical mountain, and tropical forests (SI *Appendix* Fig. S4). The classification of these forest types was based on the North American Forest map, aligned with FAO ecological zones and the CEC Terrestrials Ecological Regions of North America (45), providing representation of diverse North American forests characterized by variations in macroclimate, climate history, flora, and evolutionary lineages. For each forest type, we conducted the same SEM model to test whether the indirect effects of our three hypotheses on the relationship between species richness and basal area mirror the pattern observed across the continental scale (see goodness-of-fit for all these six SEMs in SI *Appendix* A).

## Acknowledgments

This work was done under the project P20-01233 funded by Consejería de Transformación Económica, Industria, Conocimiento y Universidades de la Junta de Andalucía and by “European Union/FEDER”. XSM is supported by “Juan de la Cierva – Formación” fellowship from the Spanish MICINN (FJC2021-047175-I). XSM, PRB, JA and JTT were funded by the Agencia Estatal de Investigación, Ministerio de Ciencia e Innovación (AEI, subproject LARGE, No PID2021-123675OB-C41). OG acknowledges financial support from provided by the Ministerio de Ciencia, Innovación y Universidades (Excelencia Europea 2023 BIOTA, No EUR2023-143472). PRB was supported by the Community of Madrid Region under the framework of the multi-year Agreement with the University of Alcalá (Stimulus to Excellence for Permanent University Professors, EPU-INV/2020/010). FRS was supported by VI PPIT-US and grants US-1381388 from Universidad de Sevilla/Junta de Andalucía/FEDER-UE and CNS2022-135839 funded by MICIU/AEI/10.13039/501100011033 and by European Union NextGenerationEU/PRTR.

## Data, Materials, and Software Availability

All data used in this study is publicly available (except for Canada NFI data which requires a use license). Code and data used to obtain the results and figures presented in this manuscript can be found https://doi.org/10.5281/zenodo.14035513. The archived data does not contain the plots of Canada NFI due to license limitations.

