## Supplementary material for "TOP-DOWN CLIMATIC PROCESSES MODULATE BIODIVERSITY-FUNCTIONING RELATIONSHIPS ACROSS NORTH AMERICAN FORESTS": SI Appendix

1. Departamento de Ciencias de la Vida, Grupo de Ecología y Restauración Forestal, Universidad de Alcalá, 28805, Alcalá de Henares, Madrid, Spain.

2. Estación Biológica de Doñana (EBD-CSIC), Sevilla, Spain.

3. Departamento de Biología Vegetal y Ecología, Universidad de Sevilla, Sevilla, Spain.

**Figure S1.** Map of national forest inventories used in our study and predictions of the relationship between basal area and species richness for each forest inventory.

**Figure S2.** Histograms of response variables in the Structural Equation Models (i.e. basal area, stem density, maximum DBH and relationship)

**Figure S3.** The relationship between mean annual temperature and annual precipitation, temperature and temperature change velocities and precipitation and temperature change velocities in our dataset.

**Figure S3.** Maps of variables stand basal area, structural and climatic variables.

**Figure S4.** Map of the distribution of the forest types evaluated.

**Figure S5.** Boxplots of the response and explanatory variables used per forest type.

**Appendix A**. Structural Equation Model details and evaluation.

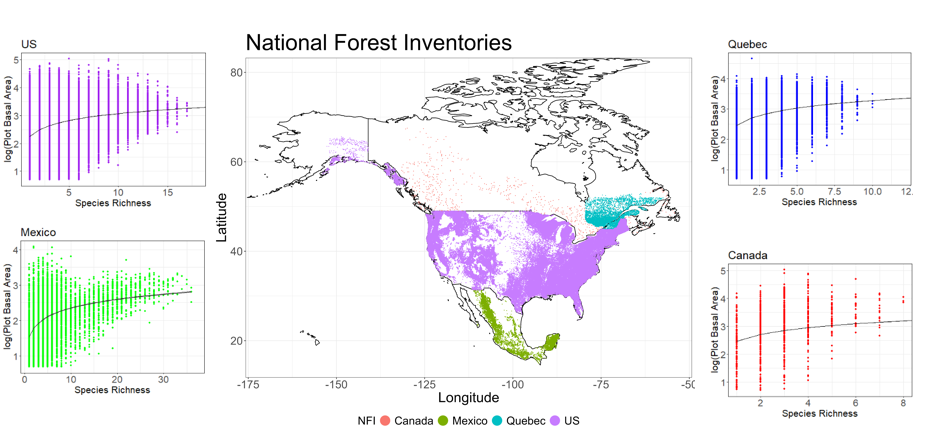

**Figure S1.** Map of the National Forest Inventories used in our study. Canada National Forest Inventory (red), Quebec Forest Inventory (blue), US National Forest Inventory (FIA, purple), and Mexico National Forest Inventory (green). Panels represent predictions of the relationship between log-transformed basal area and species richness (considering area) for each forest inventory.

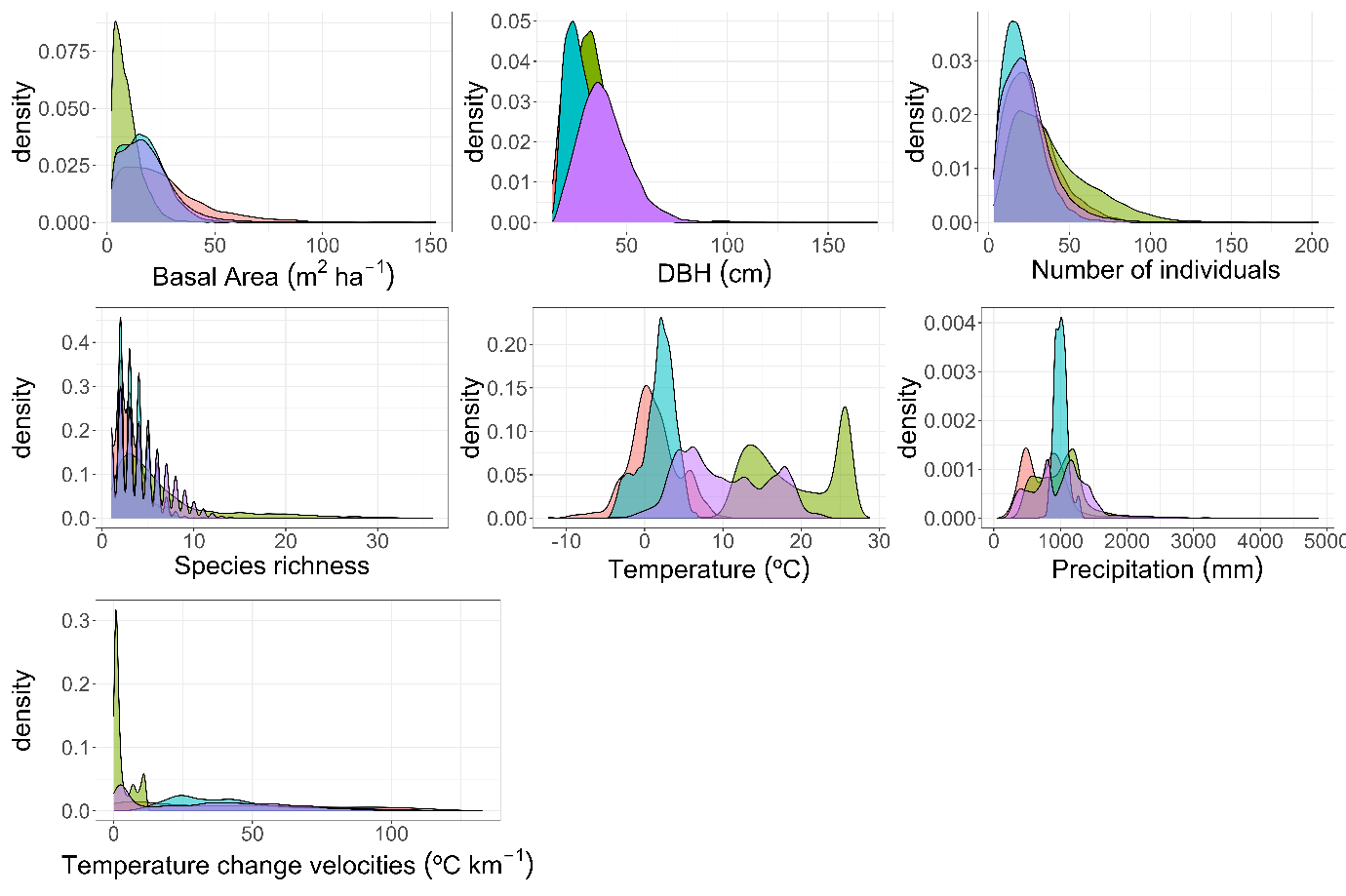

**Figure S2.** Density plots of basal area, maximum diameter (95th % percentile), number of individuals, species richness, temperature, precipitation and temperature change velocities. Color represents national forest inventory, purple = United States, red = Canada, green = Mexico, blue = Quebec.

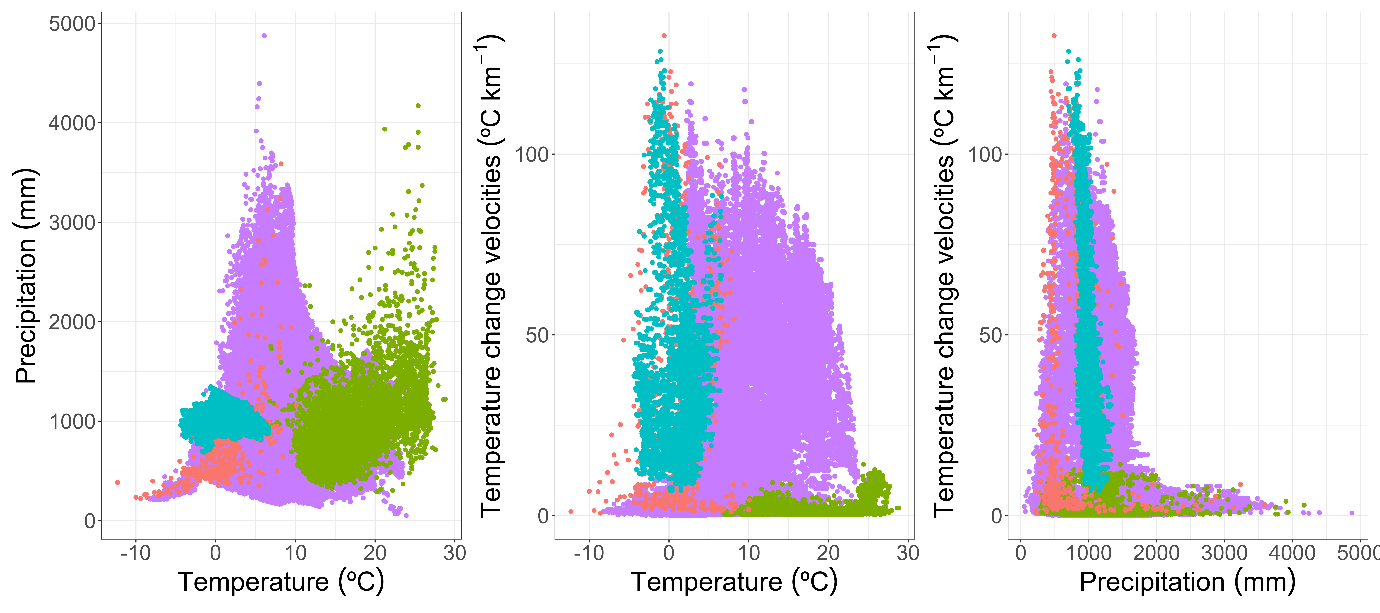

**Figure S3.** The relationship between temperature and precipitation, temperature and temperature change velocities and precipitation and temperature change velocities in our dataset. Color represents national forest inventory, purple = United States, red = Canada, green = Mexico, blue = Quebec.

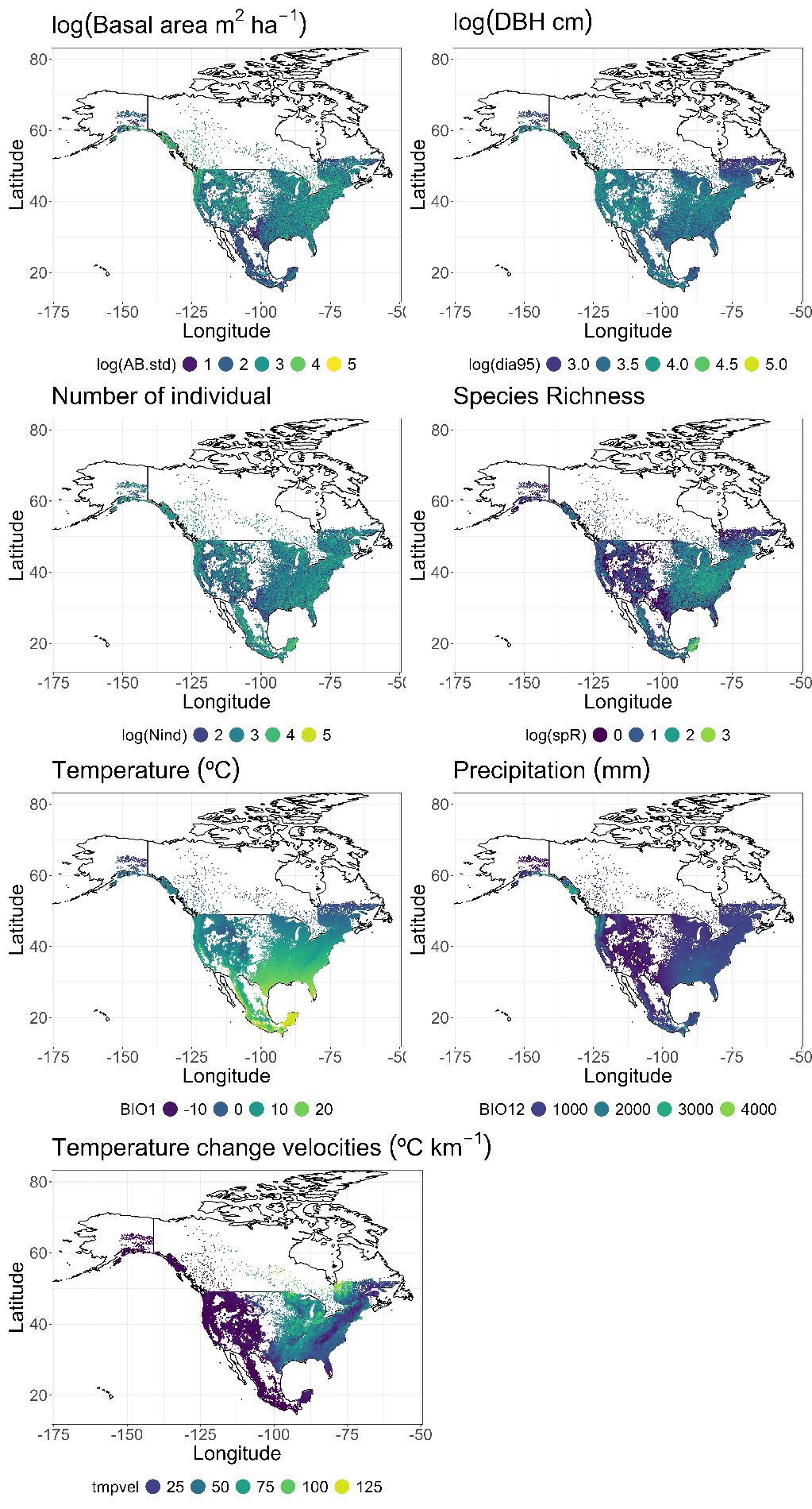

**Figure S4.** Maps of stand basal area (m2 ha-1), number of individuals, and maximum diameter (cm), temperature change velocities since the Last Glacial Maximum (ºC year-1), mean annual temperature (ºC), annual precipitation (mm).

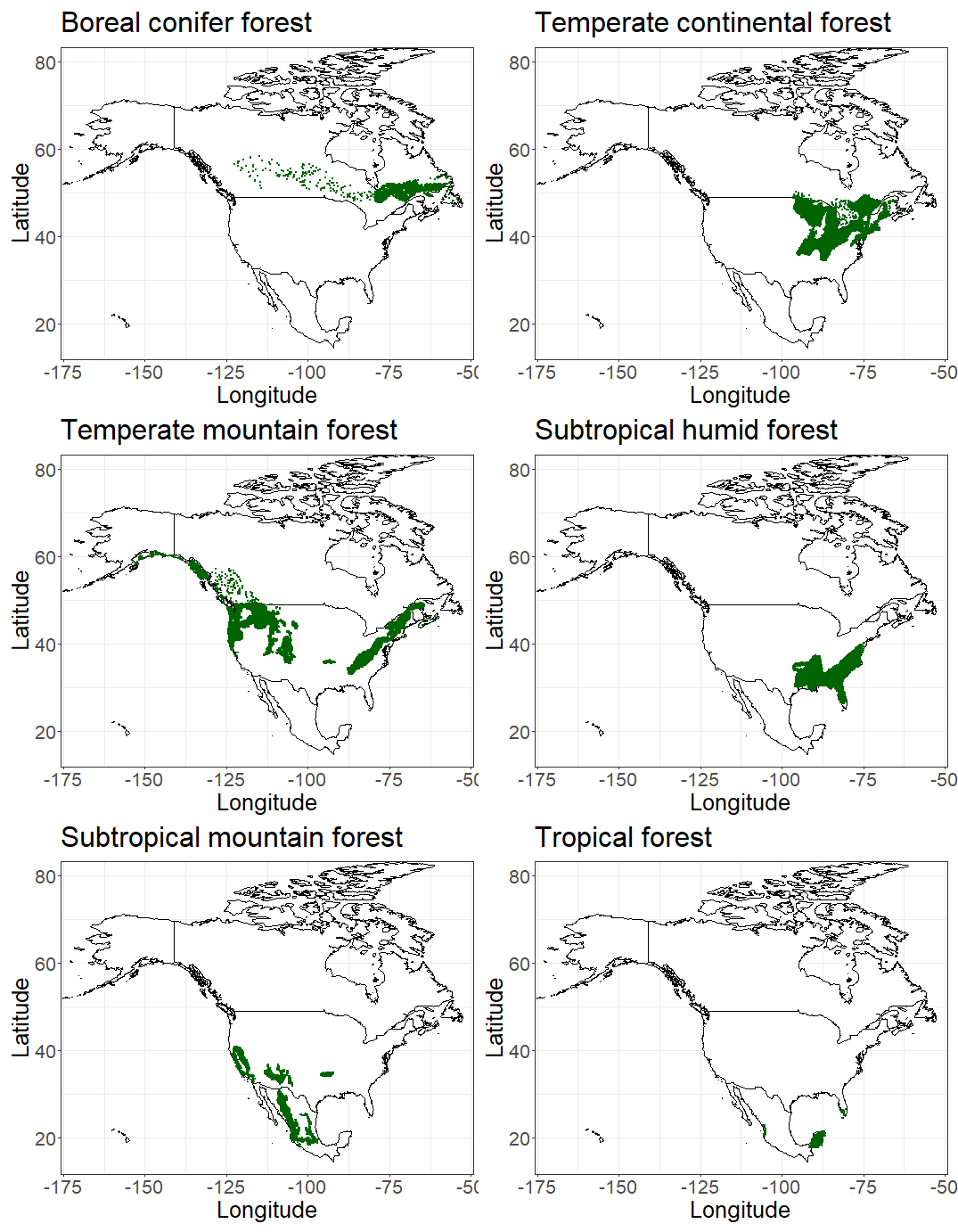

**Figure S5.** Map of the distribution of the forest types evaluated.

**Appendix A. Further details on the structural equation model**

Path justification:

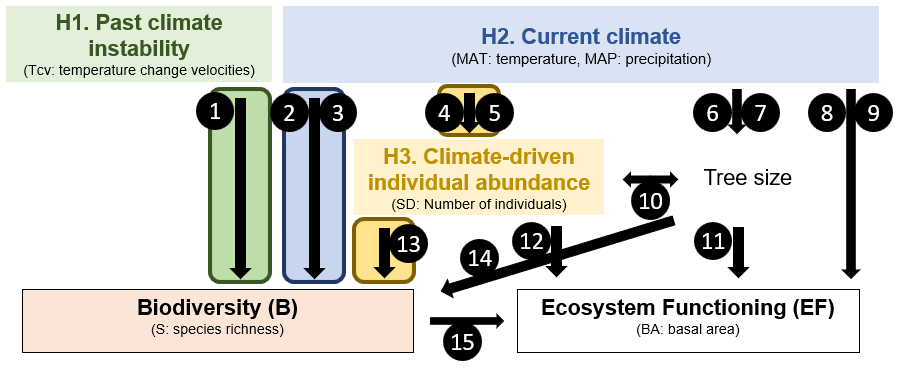

**Figure A1.** Structural Equation Model with paths numbered. In the text below there is a detailed description for the ecological justification of each of these paths.

1) **Temperature change velocity effects on species richness (path 1):** Temperature change velocities reflect the climate amplitude between the Last Glacial Maximum and the present, and more broadly, the climate change differences between glacial-interglacial periods (1, 2). It has been hypothesized that greater amplitude between these periods promotes reduced diversification and higher extinction rates, resulting in assemblages of low specialized species with large geographic ranges (3). In the case of tree diversity, temperate tree floras show legacies of past climate change that structure present-day diversity patterns (4–6). For instance it has been shown that areas with high climate change velocity present more nestedness and lower turnover of angiosperm species, indicating non-random selective processes during glacial-interglacial periods (7). These results suggest that past climate change determines current regional species pools (8) limiting the local diversity that we can observe at the plot level. However, the indirect effects of past climate change instability and biodiversity-ecosystem function patterns remain little explored (2), providing rationale for including this path in our models. **This path describes hypothesis 1 (H1) of the SEM (Figure 1), referred to as the Past Climate Instability Hypothesis**
2) **Precipitation and temperature effects on species richness paths (paths 2, 3):** Mean annual temperature and annual precipitation describe the influence of water-energy dynamics on broad-scale current diversity patterns(9) However, not all ecosystems are diversity-limited by the same environmental driver. For instance, in warm areas, water may better explain diversity patterns whereas in cold areas, temperature can be more important (9). In addition, temperature might be critical, not only because it filters species in response to the environment, but also because temperature affects metabolic rates determining mutation and speciation rates, which ultimately influence species richness (10, 11) as proposed by the “Metabolic Theory of Biodiversity”. In fact, recent work suggests the pivotal role of this theory to effectively predict latitudinal diversity gradients in tree species, although it underpredicts tropical tree diversity (12). **These two paths describe hypothesis 2 (H2) of the SEM, referred to as the Current Climate Hypothesis.**
3) **Temperature and precipitation effects on abundance, maximum tree size and stand basal area (paths 4, 5, 6, 7, 8, 9):** These three paths represent how macroclimate influences forest structure and its indirect effects on biomass accumulation. The direct paths of climate on basal area are included to control for the effects of macroclimate on biomass productivity, which is limited by temperature and water availability (13). Furthermore, previous studies have shown that temperature and precipitation have positive effects on tree density, although these effects could vary across biomes. For maximum tree size, it is also expected that precipitation and temperature will have an influence (14). However, for both tree size and number of individuals, the effects of temperature and precipitation may interact and not be straightforward, varying by forest type depending on which variable is more limiting.
4) **Covariation between maximum tree size and number of individuals (path 10):** We allowed the covariation between the number of individuals and tree size to acknowledge density dependent rules such as Yoda’s law (15). This relationship describes the effects of competition on forest structure dynamics through a self-thinning process.
5) **Maximum tree size effect on basal area (path 11):** This path allows control for the effect of forest structural diversity on basal area. Large trees have been identified as critical drivers of biomass and carbon stocks in forests, overriding climate, species diversity and other variables (16, 17). Furthermore, tree size may serve as an intermediate variable to represent an indirect effect of climate on biomass through increased productivity (18). Moreover, tree size reflects forest structure and indirectly accounts for many short-term processes crucial for biomass accumulation, such as forest management, disturbances, and stand age. While we cannot control these variables in our analyses, their influence on stand level basal area occurs through the alteration of forest structure, in other words, size class structure. Maximum tree size is positively related to stand age, indicating the absence of logging and disturbances, and therefore is a good surrogate of these variables.
6) **Number of individuals and basal area (path 12):** Following the same rationale as above, including the path for the effect of the number of individuals on basal area allows to control for the effect of forest structure on stand level basal area. Furthermore, the number of individuals may also be indicative of management, disturbance, and successional stage. Therefore, the inclusion of this path allows account for these small temporal and spatial scale processes. Furthermore, it may also show indirect effects of climate on basal areas by impeding population growth and limiting species abundance.
7) **Number of individuals effect on species richness (path 13)**: The inclusion of this path allows testing for the sampling effect on the distribution of species diversity. That is, it is likely to find plots containing more species when there are more individuals. This can be a pure probabilistic property or reflect ecological properties. For instance, the More Individual Hypothesis suggests that areas with more available energy (defined in terms of water and temperature for primary producers) support higher species abundances and, therefore, more viable populations (19, 20). **This path relates to our hypothesis 3 (H3) of the SEM (Figure 1), referred to as the Climate-driven Individual Abundance Hypothesis.**
8) **Maximum tree size and species richness (path 14):** This path is included to control for the effects of structural diversity on species richness. More complex structural forests are expected to host a greater range of tree sizes and, therefore, show a positive relationship between maximum tree size and species richness. Furthermore, the presence of larger trees is also indicative of low management and disturbance, which may be positively related to tree species richness.
9) **Species richness effect on basal area (path 15):** This path represents the widely described positive relationship between biodiversity and productivity (21), although with different strengths between forest types (22, 23).

We also included four additional paths to account for plot area in species richness, basal area, maximum diameter, and number of individuals (not shown in Figure 1 nor Fig A1).

**Structural Equation Model evaluation**

To allow for the accommodation of non-normal distributions and hierarchical structures within the model, piecewise SEM uses a local estimation method in which each endogenous variable is estimated separately (24). The local estimation method consists of the tests of direct separation (*d-sep test*), which checks for any missing relationships in the model. The d-sep test consists of three different steps. First, the creation of the *basis set,* which is the list of independence claims (the missing paths in our model that we assume have no relationship). Second, test the relationship of each independence claim. Note that if in our theoretical model we modeled a variable with certain distribution, all independent claims to this variable will be tested using such distribution (e.g., number of individuals as negative binomial distribution). A p-value is obtained from this relationship, if the p-values > 0.05 we fail to reject the null hypothesis (variables are conditionally independent) and thus support the exclusion of the path. Third, there is an evaluation of the overall model fit. The probabilities *p_i_* of each of the *k* independent claims are obtained, which are then combined into the Fisher’s *C* statistic:

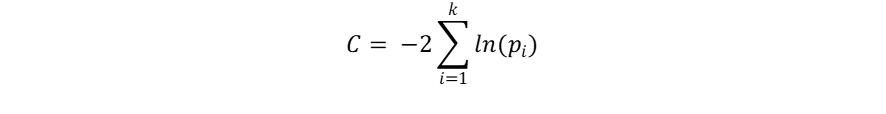

Where *k* is the number of independent claims, *i* the independence claim, and *p* the p-value of the independent claim. If *Fisher C* is > 0.05 we cannot reject the null hypothesis (i.e. the model represents well our data). Fisher’s *C* statistic is used to obtain a value of Akaike’s Information Criterion (AIC) as: AIC = C + 2K, where *K* is the likelihood of degrees of freedom.

Although our SEM is theory-based, it is possible for independence claims lacking biological or ecological relevance to be supported. In such cases, the model can be qualitatively evaluated with coefficients of determination from each submodel. Removing independence claims can be a subjective decision, as can the inclusion of relationships in the model after seeing the improvement on the model fit. Given that all variables in our dataset are statistically related and our sample size is large enough to detect even minor and potentially irrelevant relationships, we consistently reject the null hypothesis of our independent claims (i.e., that there is no relationship between the variables) and our global model evaluation (i.e., that the model correctly represents the data). Therefore, we concluded that the most appropriate approach is to estimate the model fit for each model used to construct the SEM (see 25).

In all continental models, plot area was included as a fixed effect to account for area effects and possible differences between forest inventories. Therefore, despite being common for all submodels specified below, we do not mention it again.

**SEM Submodels**

**Submodel 1. Stand basal area.** Basal area was log-transformed with a normal error distribution as a function of species richness, number of individuals, log transformed tree size, mean annual temperature and annual precipitation. All response variables except the number of individuals were scaled and centered.

**Submodel 2. Stem density.** To model stem density, we fitted a generalised linear model with a negative binomial distribution. We included as fixed effects mean annual temperature and annual precipitation, whereas tree size was not included due to its covariation with number of individuals.

**Submodel 3. Tree size.** To model maximum tree diameter within each plot we fitted a general linear model of log-transformed tree size with a normal error distribution. We included as fixed effects mean annual temperature and annual precipitation.

**Submodel 4. Species richness.** To model species richness we fitted a generalised linear effect model with a Poisson distribution. We included as predictors mean annual temperature and annual precipitation, temperature change velocities, stem density and tree size.

**Table A1.** R2 for SEM submodels, where the response variable is stand basal (m^2^ ha^-1^), Number of individuals (No. trees), tree size (cm), species richness (No. tree species).

| **Model response variable** | **R2** |
| --- | --- |
| Stand basal (m2 ha-1) | 0.86 |
| Number of individuals | 0.13 |
| Tree size | 0.1 |
| Species richness | 0.48 |

**Table A2.** Marginal and conditional R2 for SEM models per forest type, where the response variable is stand basal (m^2^ ha^-1^), stem density (No. trees), tree size (cm), species richness (No. tree species). We quantified marginal and conditional R2.

| Forest types | Stand basal area | Stem density | Tree size | Species richness |
| --- | --- | --- | --- | --- |
| **Tropical** | 0.91 | 0.08 | 0.11 | 0.93 |
| **Subtropical mountain** | 0.85 | 0.3 | 0.1 | 0.42 |
| **Subtropical humid** | 0.86 | 0.01 | 0.01 | 0.49 |
| **Temperate continental** | 0.86 | 0.09 | 0.12 | 0.42 |
| **Temperate mountain** | 0.85 | 0.07 | 0.06 | 0.6 |
| **Boreal** | 0.9 | 0.01 | 0.07 | 0.37 |

Model residuals were checked using the *DHARMa* R package (26). In the following figures we show the residual plots for all submodels in each SEM (all North America and the evaluated forest types).

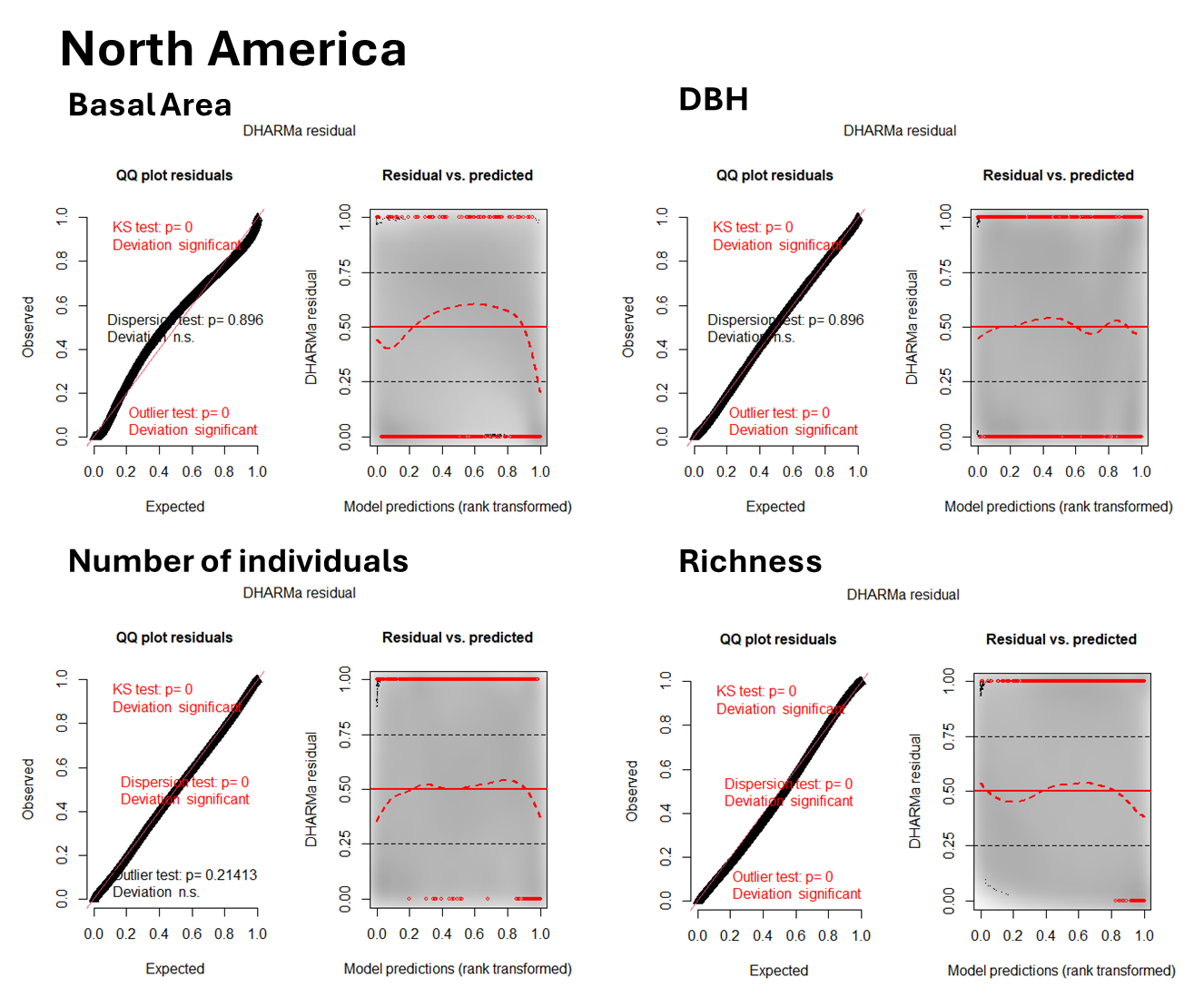

**Figure A1.** qqplots and plots of residuals against predicted values generated with DARMa r package (26) for SEM submodels of all North America data. Response variables were basal area (submodel 1), 95th percentile plot DBH (submodel 2), number of individuals (submodel 3) and species richness (submodel 4).

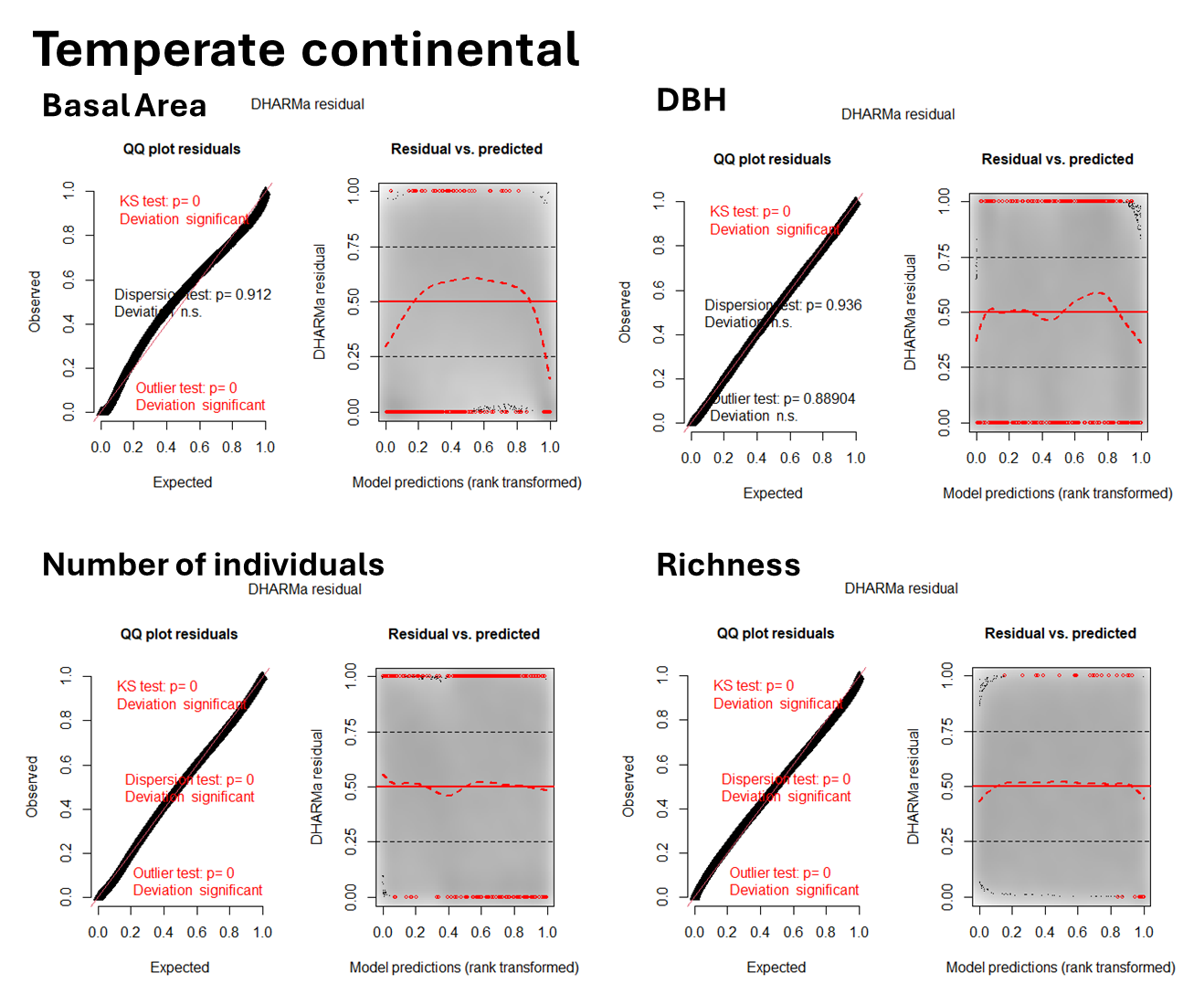

**Figure A2.** qqplots and plots of residuals against predicted values generated with DARMa r package (26) for SEM submodels of temperate continental forest data. Response variables were basal area (submodel 1), 95th percentile plot DBH (submodel 2), number of individuals (submodel 3) and species richness (submodel 4).

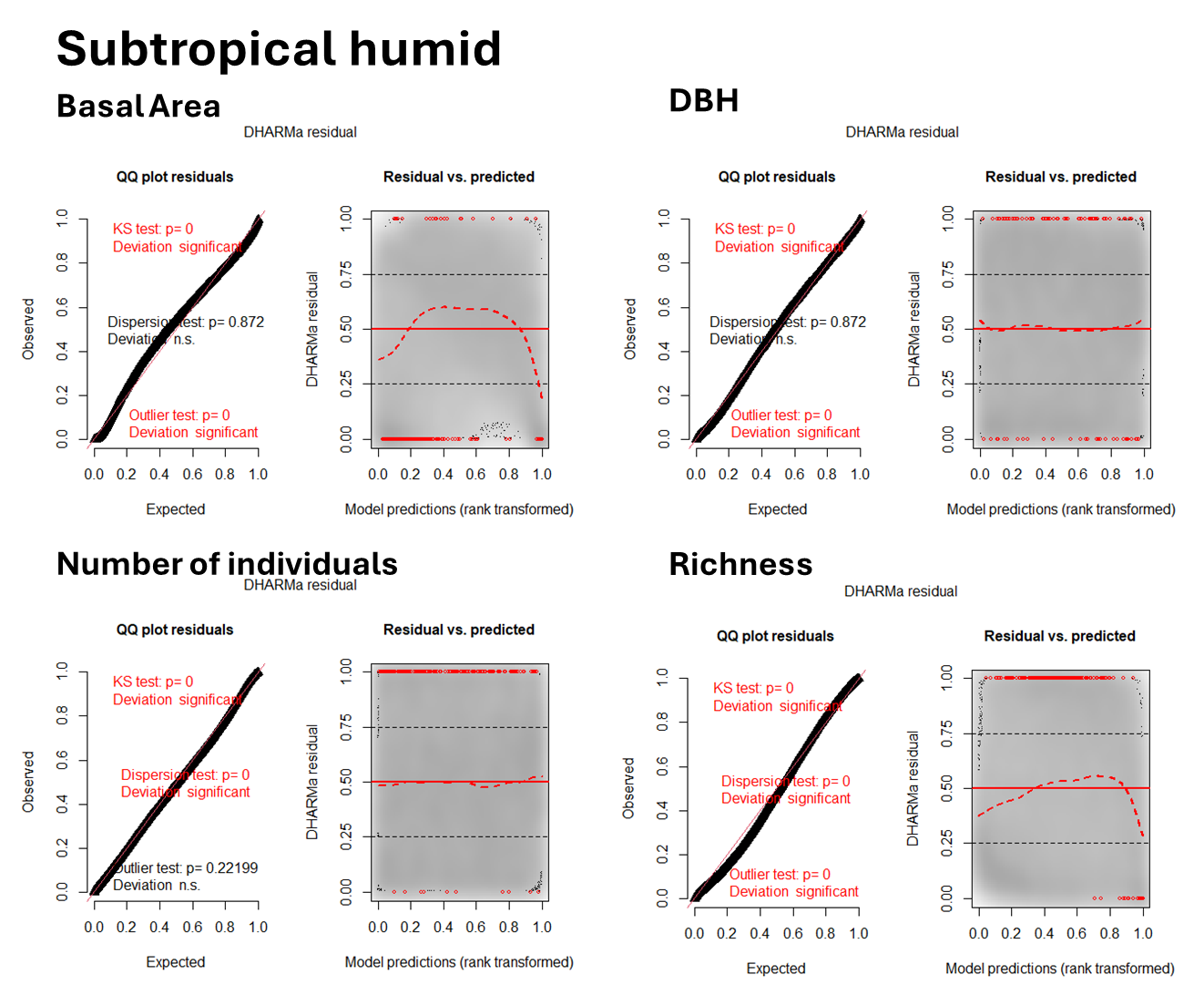

**Figure A3.** qqplots and plots of residuals against predicted values generated with DARMa r package (26) for SEM submodels of subtropical humid forest data. Response variables were basal area (submodel 1), 95th percentile plot DBH (submodel 2), number of individuals (submodel 3) and species richness (submodel 4).

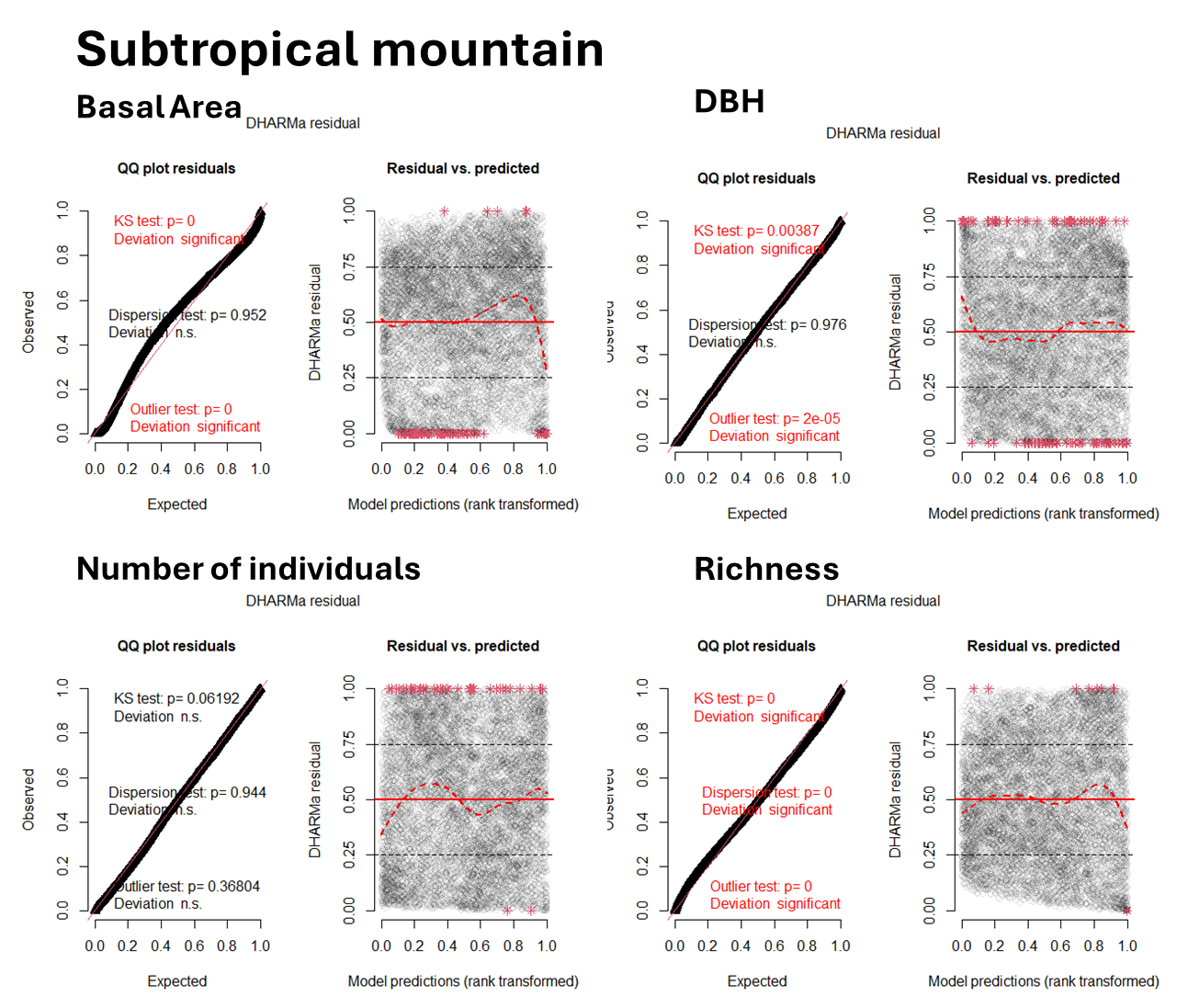

**Figure A4.** qqplots and plots of residuals against predicted values generated with DARMa r package (26) for SEM submodels of subtropical mountain forest data. Response variables were basal area (submodel 1), 95th percentile plot DBH (submodel 2), number of individuals (submodel 3) and species richness (submodel 4).

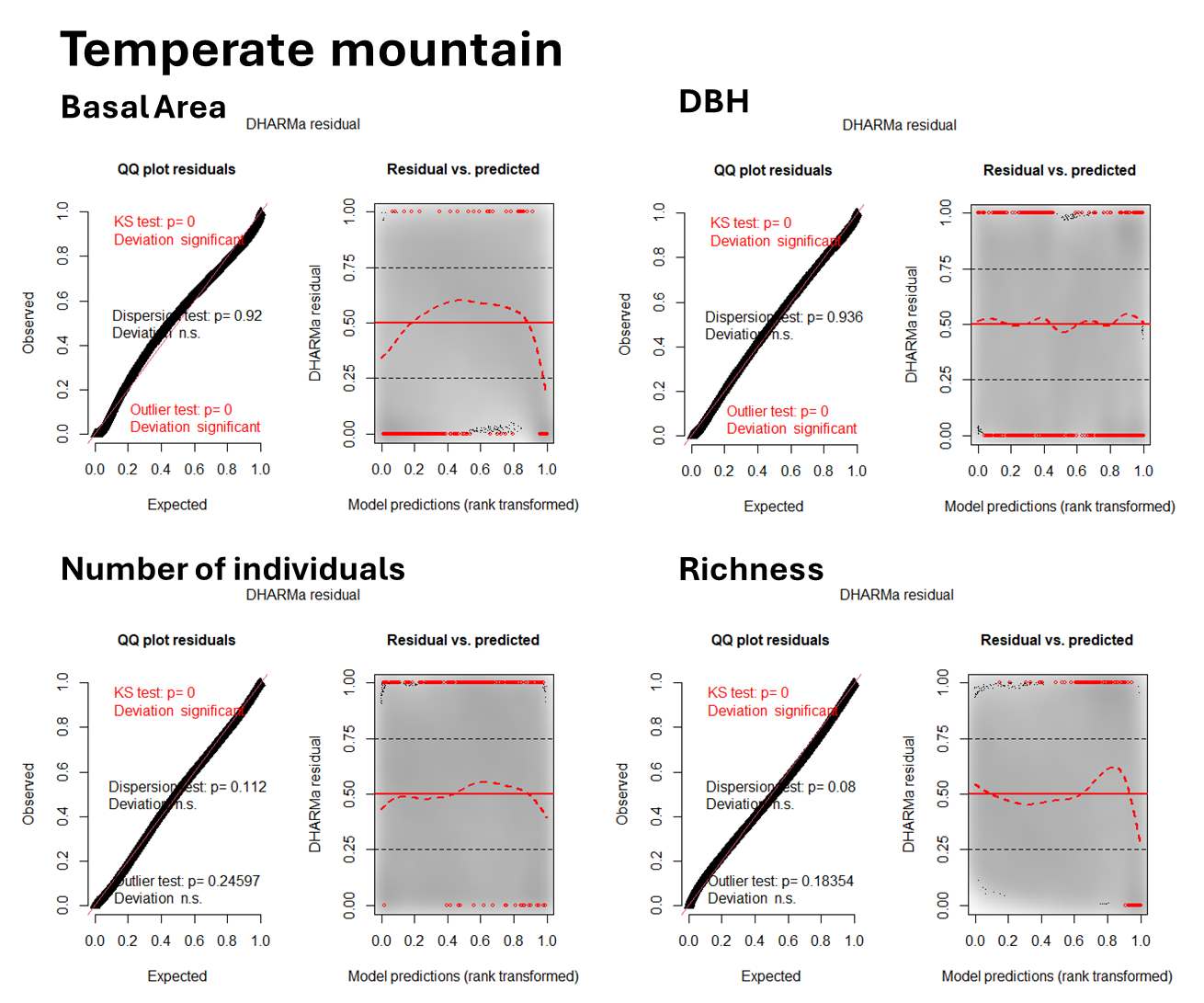

**Figure A5.** qqplots and plots of residuals against predicted values generated with DARMa r package (26) for SEM submodels of temperate mountain forest data. Response variables were basal area (submodel 1), 95th percentile plot DBH (submodel 2), number of individuals (submodel 3) and species richness (submodel 4).

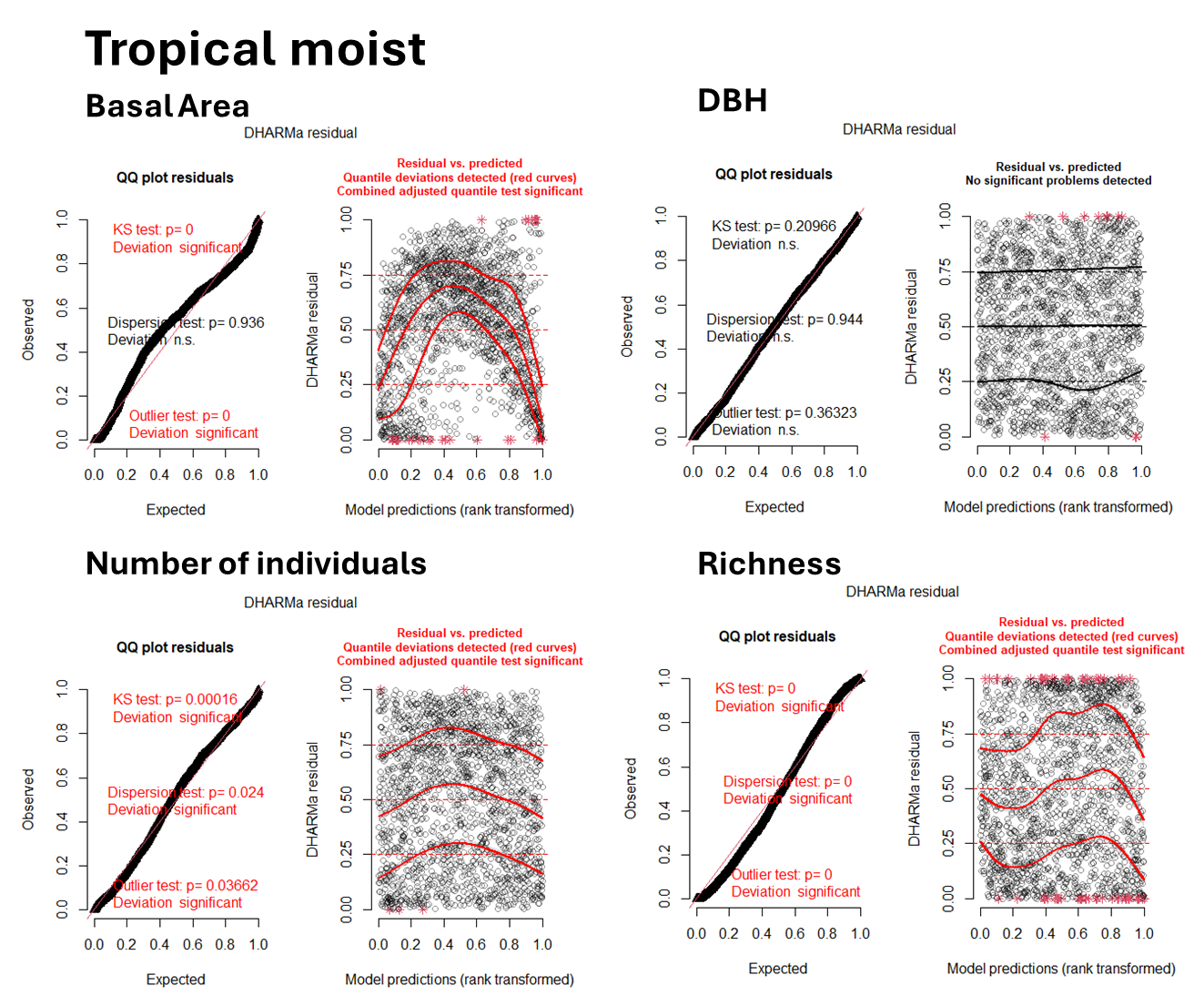

**Figure A6.** qqplots and plots of residuals against predicted values generated with DARMa r package (26) for SEM submodels of tropical moist forest data. Response variables were basal area (submodel 1), 95th percentile plot DBH (submodel 2), number of individuals (submodel 3) and species richness (submodel 4).

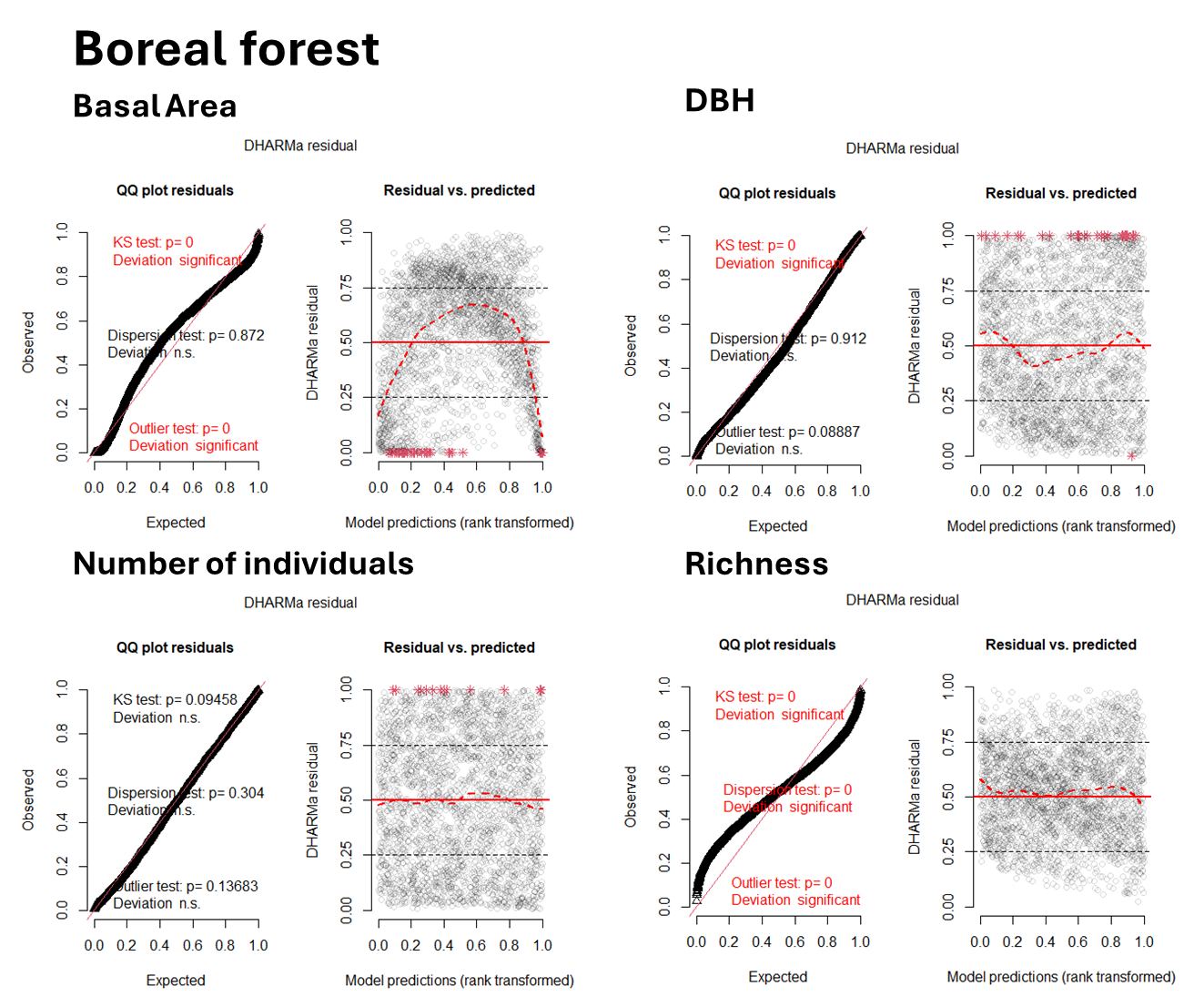

**Figure A7.** qqplots and plots of residuals against predicted values generated with DARMa r package (26) for SEM submodels of boreal conifer forest data. Response variables were basal area (submodel 1), 95th percentile plot DBH (submodel 2), number of individuals (submodel 3) and species richness (submodel 4).

26. DHARMa: residual diagnostics for hierarchical (multi-level/mixed) regression models. R package version 0.4. 6
